# Autoimmune non-coding variants perturb transcription factor–cofactor complex assembly linked to enhancer activity

**DOI:** 10.64898/2026.05.20.726379

**Authors:** Maryam Dashtiahangar, Trevor Siggers

## Abstract

Most autoimmune disease-associated variants lie in non-coding regions, but the molecular mechanisms linking these variants to gene regulation remain poorly understood. A major unresolved challenge is to determine how disease alleles alter transcription factor (TF) binding, cofactor (COF) recruitment, and enhancer activity at scale. Here, we used the CASCADE method to profile differential binding of five TFs and ten COFs to 2,901 autoimmune disease-associated variants in Jurkat T cells, identifying 516 binding-modulating variants. Variants impacting binding were enriched among MPRA-defined expression-modulating variants and were strongly concordant with allele-specific reporter expression, linking altered TF/COF recruitment to enhancer activity. A majority of variants perturb binding of five major TF families — ETS, RUNX, SP/KLF, OVOL/MYBL, and bHLH — all of which have established roles in T cell biology. Notably, we find that ETS and RUNX factor binding is enriched at different variant functional classes, suggesting that they act through distinct regulatory mechanisms at disease loci. We describe allele-dependent regulator “switching” at several loci, where distinct complexes are found at reference and variants alleles, and we identify a recurrent regulatory module involving FOXM1 and the cofactors TIP60, BRD4, NCOA3, and NCOA1 assembling on ETS sites that tracks with gene expression. Together, this integrated biochemical and functional framework prioritizes autoimmune disease-associated variants by linking allele-specific TF/COF binding mechanisms to enhancer activity.

## INTRODUCTION

Most genetic variants associated with autoimmune disease lie outside protein-coding regions, implicating changes in gene regulation as a major mechanism of disease risk^1,2^. Genome-wide association studies (GWAS) have identified thousands of non-coding variants (NCVs) linked to immune-mediated diseases, including type 1 diabetes^3^, inflammatory bowel disease^4^, rheumatoid arthritis^5^, psoriasis^6^, and multiple sclerosis^7,8^. However, identifying the causal variants within disease-associated loci remains challenging because GWAS signals often contain many correlated variants in linkage disequilibrium (LD). Determining how these variants alter regulatory mechanisms is also difficult because NCVs can affect transcription factor (TF) binding, chromatin state, enhancer activity, and gene expression in a cell-type-specific manner. Thus, mechanistic interpretation requires approaches that connect individual alleles to their molecular regulatory consequences.

Several experimental and computational strategies have been developed to identify putative functional NCVs. MPRAs identify variants that alter enhancer or promoter activity^9–15^, while CRISPR-based perturbation screens can link regulatory elements to target-gene expression in their native genomic context^16–19^. Chromatin-accessibility^2,20–22^ and histone-mark profiling^23–27^ identify variants located within active regulatory regions, and eQTL^28–31^ and caQTL/chromating QTL^32–36^ mapping connect genetic variation to changes in gene expression or chromatin accessibility. In parallel, computational motif-disruption approaches^37–39^ and experimental TF-binding assays, including EMSAs^40,41^, SELEX-based variant-binding assays^42,43^, and others^44,45^

,can predict or directly measure allele-specific effects on TF binding. However, defining the mechanism of a regulatory variants requires identifying not only the TFs affected by each allele, but also the regulatory cofactors (COFs) they recruit. COFs are central regulators of enhancer function^46–48^ that are recruited by DNA-bound TFs and impact gene expression via diverse mechanisms, including histone modification and chromatin remodeling. Thus, disease-associated variants may alter gene regulation by changing TF binding directly or by disrupting broader TF– COF regulatory complexes. Because these TF-COF complexes depend on both DNA sequence, protein-protein interactions, and cellular context, they cannot be fully inferred from motif disruption or TF binding alone. A scalable experimental approach is therefore needed to measure how disease-associated alleles alter both TF binding and COF recruitment in a cell-type- and state-specific manner.

CASCADE (Customizable Approach to Survey Complex Assembly at DNA Elements) provides a framework to address this gap by profiling TF and COF binding to variant-containing DNA sequences using nuclear extracts^49,50^. CASCADE uses protein-binding microarrays (PBMs)^49,51,52^ to quantify binding of TFs and COFs present in nuclear extracts to DNA probes (Fig. 1a). Here, we used CASCADE to allele-specific recruitment of select TFs and COFs to 2,901 autoimmune disease-associated NCVs previously characterized by massively-parallel reporter assay (MPRA) in Jurkat T cells^9^. We identified 516 bmVars across the autoimmune variant set. These bmVars were enriched among MPRA expression-modulating variants (emVars) and showed strong concordance between the direction of binding changes and reporter expression changes, supporting a direct link between altered regulatory complex assembly and enhancer activity. CASCADE also identified bmVars among MPRA-active putative cis-regulatory elements that lacked allelic expression effects (pCREs), revealing pCRE/bmVars with regulatory binding changes that may be missed when variants are prioritized based only on allele-specific reporter expression. Motif analysis implicated several major TF families, including ETS, RUNX, SP/KLF, OVOL_MYBL, and bHLH factors, with distinct enrichment patterns between emVar/bmVars and pCRE/bmVars. At individual loci, CASCADE revealed allele-dependent switching between TF-family binding sites and identified a recurrent ETS-associated coactivator module whose allele-dependent recruitment was linked to enhancer activity. Comparison with motifbreakR^37^ showed that sequence-based predictions often over-nominate disrupted TF families and fail to capture context-dependent binding changes, particularly at RUNX-associated loci.

**Figure 1.**
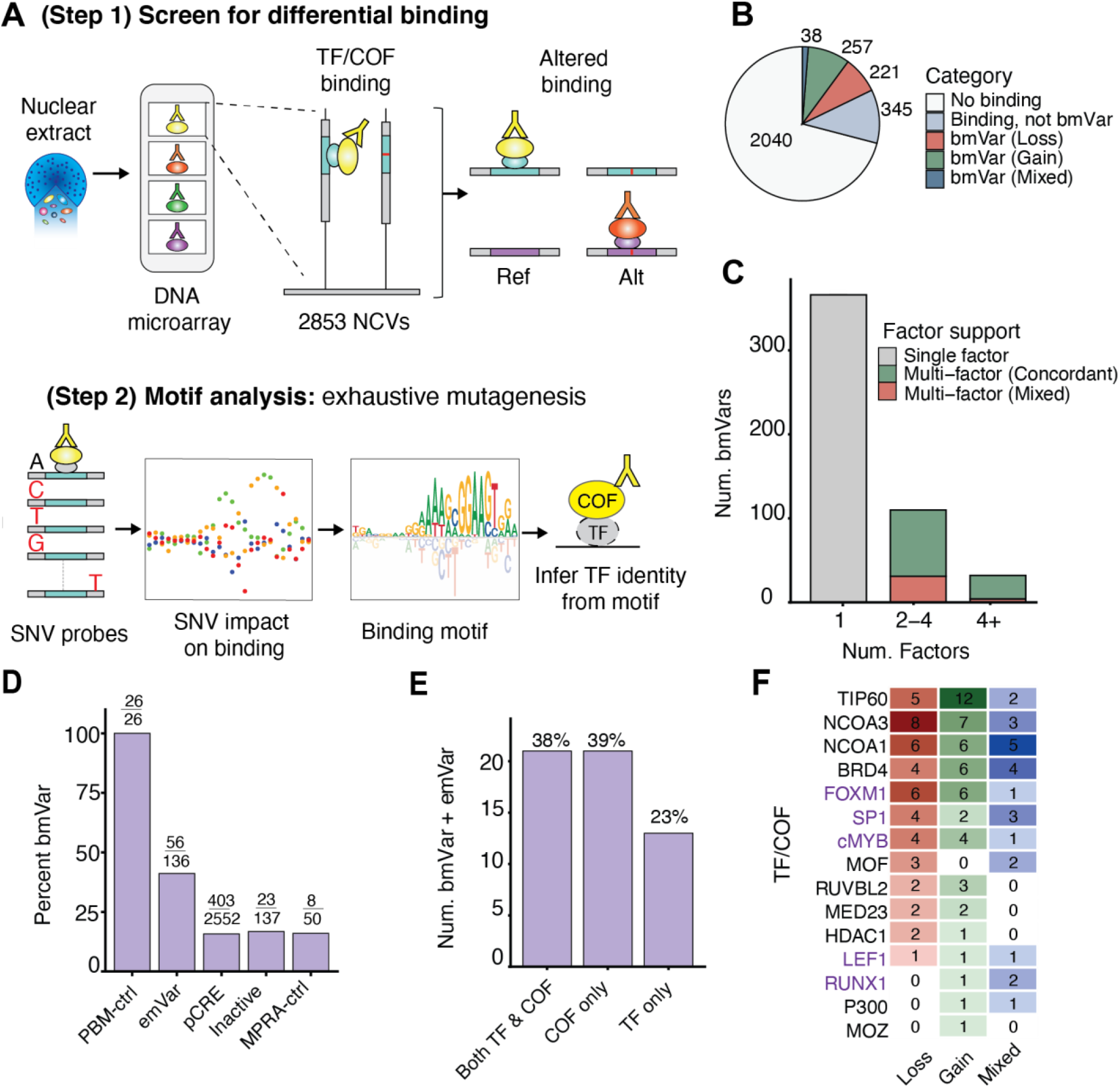
CASCADE profiling of TF and COF binding at autoimmune disease-associated variants. **(a)** Schematic of the CASCADE two-step approach to profile factor binding to non-coding variants (NCVs). In Step 1, DNA-binding of factors from cell nuclear extracts are profiled against reference (Ref) and alternate (Alt) alleles on a DNA microarray. In Step 2, exhaustive single-nucleotide variant (SNV) mutagenesis arrays are used to determine factor recruitment motifs at prioritized loci, which are compared to TF motif databases to infer the identity of TF families whose binding is disrupted by each variant. **(b)** Classification of variants based on Step 1 results: binding-modulating variants (bmVars) showing gain, loss, or mixed effects; no detectable binding (No binding); detectable but not different between alleles (not bmVar). **(c)** Number of factors supporting each bmVar. Multi-factor bmVars are classified as directionally concordant when all supporting factors change in the same direction, or mixed when factors show opposing directions. **(d)** Variant fraction classified as bmVars for MPRA-defined variant categories and CASCADE positive controls. **(e)** Distribution of bmVar/emVars based on whether differential binding affected TFs only, COFs only, or both TFs and COFs. **(f)** Factor-level summary of loss, gain, and mixed binding effects among bmVar/emVars.

To our knowledge, this is the first study to integrate high-throughput biochemical profiling of TF and COF binding with MPRA-based functional measurements across thousands of autoimmune disease–associated NCVs. This combined framework identifies high-priority variants supported by both molecular mechanism and enhancer activity, while also revealing MPRA-active variants with binding changes that are not captured by allelic expression effects alone. Importantly, our results highlight COF recruitment as an informative and experimentally accessible layer of variant mechanism, extending functional variant interpretation beyond TF binding alone.

## RESULTS

### Profiling TF and COF binding at autoimmune disease-linked NCVs in T cells

To investigate the regulatory mechanisms of autoimmune disease-associated NCVs, we used CASCADE to profile differential binding of TFs and COFs to 2,901 variants previously characterized by MPRA in Jurkat T cells^9^. These variants were linked to five autoimmune diseases: type 1 diabetes (T1D), inflammatory bowel disease (IBD), rheumatoid arthritis (RA), psoriasis, and multiple sclerosis (MS). To screen a large numbers of variants, we employ a hierarchical two-step approach^49^. In Step 1 (Screen), TF and COF binding to paired Ref/Alt allele probes is screened to identify variants that lead to significant differential binding — we term these binding-modulating variants (bmVars) (Fig. 1a, Step 1). In Step 2 (Motif analysis), a second microarray is used to determine CASCADE-based TF/COF motifs at each bmVar locus by profiling binding to all single-nucleotide variants (SNV) probes (Fig. 1a, Step 2). For TF experiments, Step 2 identifies the DNA motifs bound by the profiled TF; however, as COFs do not bind DNA directly, motifs identified in COF experiments are used to infer the TFs recruiting them to that loci. TF inference is done by comparing COF motifs to TF motif databases. As many TFs recognize similar DNA-binding motifs, CASCADE assigns TF identities at the level of ‘TF motif clusters’ rather than individual TFs^49,53^. Differential TF binding identifies variants that directly alter recruitment of DNA-binding regulators, while differential COF binding provides complementary information about additional layers of regulation, including how variants may impact chromatin state through altered recruitment of epigenetic regulators. Further, because COFs interact with a wide range of TFs, profiling a single COF can provides readout of multiple TF–COF complexes across variant loci, without requiring prior knowledge of the TFs bound at each site.

Using nuclear extracts from resting Jurkat T cells, we profiled five TFs (SP1, FOXM1, cMYB, RUNX1, LEF1) implicated in T cell-mediated autoimmunity and ten COFs spanning diverse functional classes, including histone acetyltransferases (TIP60, P300, NCOA1/SRC1, NCOA3/SRC3), a deacetylase (HDAC1), a bromodomain scaffold (BRD4), and a Mediator subunit (MED23) (Supplementary Table 1). The variant library comprised Ref/Alt nucleotide pairs for the 2901 autoimmune disease-associated variants described above: 136 expression-modulating variants (emVars); 2,552 putative cis-regulatory elements (pCREs) that showed expression by MPRA but not allele-specific expression; 137 variants in tight linkage disequilibrium (LD) with lead SNPs that were inactive by MPRA (Hi LD, not active); 50 variants distal to lead SNPs that were inactive by MPRA (Low LD, not active). Also included were 26 Ref/Alt pairs previously shown by CASCADE to exhibit allele-specific binding (CASCADE ctrl)^49^ (Methods).

### Variants perturb TF and COF binding at hundreds of loci

We identified statistically significant differential binding for at least one TF or COF for 516 of 2,901 variants tested (17.8%) (Methods; Supplementary Table 3, Sheet 1). We refer to variants that impact binding as binding-modulating variants (bmVars). Among bmVars, 49.8% (257/516) were gain-of-binding events, 42.8% (221/516) were loss-of-binding events, and 7.3% (38/516) showed “mixed effects” where different factors exhibited opposite perturbation directions. An additional 345/2,901 variants (11.8%) showed detectable TF or COF binding that was not significantly perturbed (Fig. 1b). 29% (150/516) of bmVars altered the binding of multiple factors, and among these multi-factor bmVars, 77.4% (112/150) showed complete directional concordance, with all supporting factors changing in the same direction, suggesting coordinated effects on regulatory complex assembly (Fig. 1c). In contrast, discordant variants suggest scenarios in which TF/COF binding is switching between different complexes (discussed further below).

### Differential regulatory binding shows concordant changes with expression

To determine whether variants that perturb TF/COF binding also affect reporter gene expression, we compared bmVars with emVars identified by MPRA^9^. Of 516 bmVars, 56 (10.8%) were annotated as emVars, representing a higher bmVar rate among emVars (41.1%) than for MPRA-negative controls and inactive elements (Fig. 1d). Among joint bmVar/emVars, 39% (22/56) perturbed binding of COFs alone and 23% (13/56) perturbed TF binding alone (Fig. 1e). Although this difference may partly reflect the larger number of COFs profiled (10 vs. 5 TFs), it demonstrates that COF profiling captures regulatory variants beyond those detected by the TFs included in our panel. EmVars most frequently perturbed the cofactors TIP60, NCOA3, NCOA1, and BRD4 (Fig. 1f). Individual bmVar/emVar profiles revealed coordinated TF/COF recruitment patterns aligned with expression changes (Fig. 2a). Binding changes in bmVar/emVars showed strong concordance with the direction of gene expression change (77.8%), significantly exceeding random expectation (50%, 10,000 permutations, p < 0.0001) and exceeding the concordance rate seen with pCREs and inactive elements lacking significant allelic expression differences (p = 0.002, Fisher’s exact test; Fig. 2b). BmVar/emVars also showed greater multi-factor support, with 40% affecting multiple TF/COF factors compared to 22% among bmVars that were not emVars (Fig. 2c).

**Figure 2.**
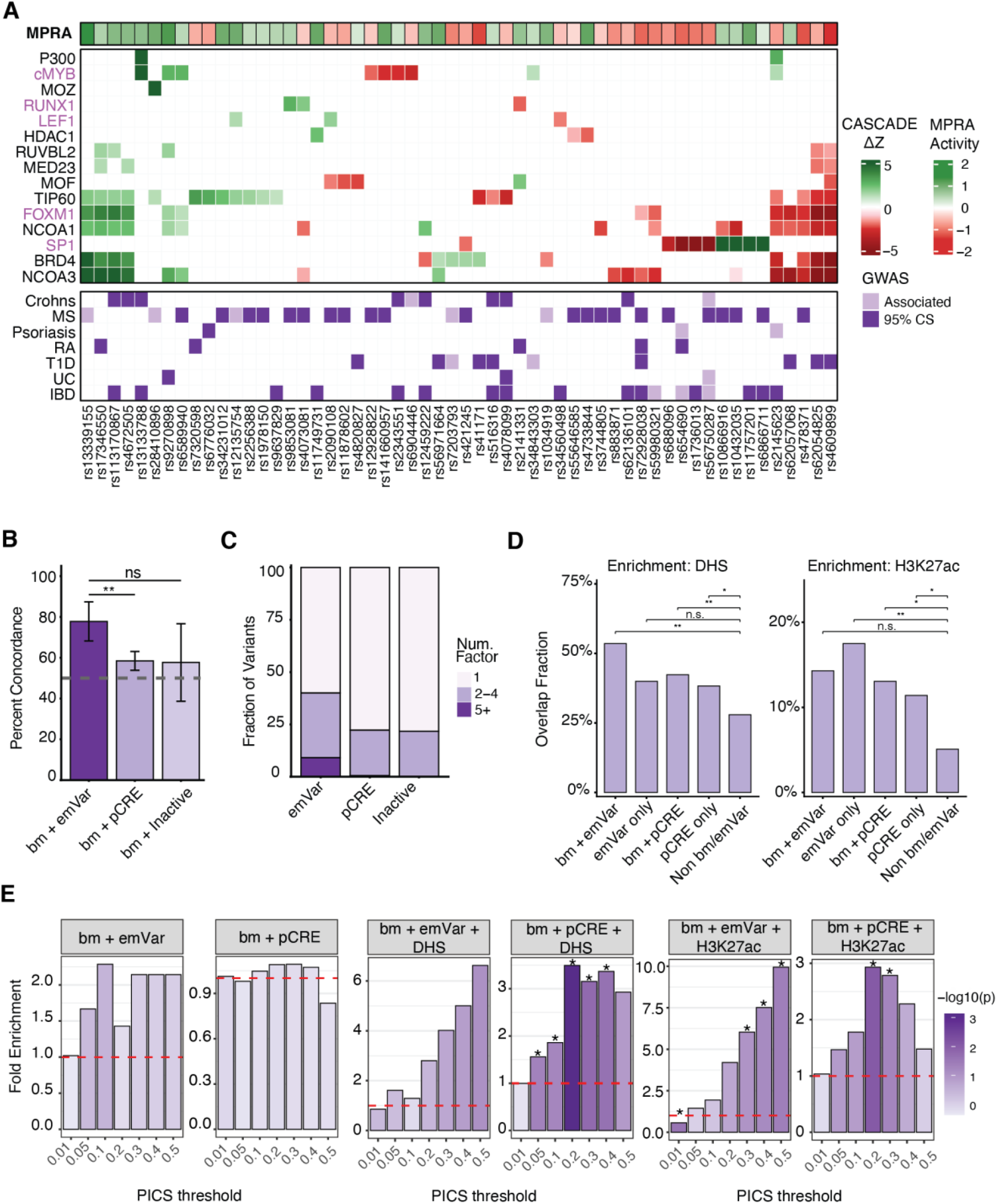
CASCADE binding changes are concordant with MPRA expression effects and enriched in functional regulatory annotations. **(a)** Summary of bmVar/emVar impact on binding, expression, and disease association. Top: MPRA reporter gene activity (green, upregulation; red, downregulation). Middle: CASCADE differential binding quantified by Δz-scores (green, binding gain; red, binding loss). TF names are in purple; COF names are in black. Bottom: GWAS disease association indicating when each variant is within the 95% credible set for each disease (dark purple) or has a GWAS association but is not in the credible set (light purple). Variants labeled by rsID. **(b)** Binding-expression concordance rates for bmVar/emVars, bmVar/pCREs, and bmVar/inactive elements. Concordance is defined as gain of binding paired with increased MPRA expression, or loss of binding paired with decreased expression. Error bars indicate 95% confidence intervals. Significance assessed by Fisher’s exact test; **p < 0.01, n.s. = not significant. **(c)** Multi-factor support across MPRA-defined variant categories. Stacked bars show the proportion of bmVars supported by 1, 2–4, or 5+ TF/COF factors among emVars, pCREs, and inactive elements. **(d)** Proportion of variants overlapping T cell DNase I hypersensitive sites (DHS, left) and H3K27ac ChIP-seq peaks (right) across five variant categories: bmVar/emVars (bm + emVar), non-bmVar emVars (emVar only), bmVar/pCREs (bm + pCRE), non-bmVar pCREs (pCRE only), and non-bmVar/MPRA-inactive variants (Non bm/emVar). Significance was assessed by Fisher’s exact test comparing each category to the non-bmVar/MPRA-inactive baseline; *p < 0.05, **p < 0.01, n.s. = not significant. **(e)** Fold enrichment of elevated PIP fine-mapping scores at multiple posterior inclusion probability (PIP) thresholds for six comparisons. For bmVar/emVar and bmVar/pCRE comparisons, fold enrichment was calculated relative to the non-bmVar category. For comparisons that also include DHS or H3K27ac overlap, fold enrichment was calculated relative to all other tested variants not in the indicated bmVar/chromatin-overlap category. Bar shading indicates statistical significance (−log10(p), Fisher’s exact test). Red dashed line indicates fold enrichment of 1 (no enrichment).

To further assess the functional relevance of bmVars, we examined their overlap with T cell DNase I hypersensitive sites (DHS), H3K27ac ChIP-seq peaks, and fine-mapping posterior inclusion probability (PIP) scores from PICS (Method) that estimate the probability that a variant is causal at a GWAS locus. As emVars were originally defined by MPRA, they are already expected to be enriched for functional regulatory annotations, including DHS, H3K27ac, and high PIP scores relative to other MPRA-tested variants. This sets a high baseline for detecting additional enrichment from bmVar status. Still, bmVar/emVar showed the highest DHS overlap and were significantly enriched in T cell DHS regions compared to inactive baseline (Not bm/emVar) (p = 0.0013, Fisher’s exact test), while H3K27ac overlap showed a similar trend that did not reach significance (p = 0.068) (Fig. 2d). bmVar status also showed enrichment for elevated PIP scores, reaching significance at multiple PIP thresholds when combined DHS or H3K27ac regions (p < 0.05, Fisher’s exact test) (Fig. 2e), suggesting that bmVar status captures functional binding perturbations specifically within open chromatin. Together, these findings demonstrate that CASCADE identifies regulatory variants that perturb TF–COF complex assembly, and that variants with concordant binding and expression changes represent a high-priority subset for further mechanistic characterization.

### NCVs predominantly impact binding for five TF families

To examine the TF families impacted by disease-associated NCVs, we performed CASCADE motif analysis (Step 2) on 56 emVar/bmVars and 113 pCRE/bmVars for the same 15 factors profiled in Step 1. Our approach defines a binding motif for each profiled factor at each locus. We identified motifs matching known TFs (adjusted p < 0.01) at 36 emVar/bmVar loci (65%) and 63 pCRE/bmVar loci (56%) and assigned them to TF motif clusters based on shared TF binding similarity (Method, Supplementary Table 2, Supplementary table 3, Sheet 2; Supplementary Table 4). For most variants with an identified motif (78/99, 79%), only one profiled TF or COF yielded a motif (Fig. 3a). At the remaining 21 loci, motifs were identified for multiple factors. Among these multi-factor motif loci, 57% (12/21) showed the same motif cluster across all factors, suggesting that multiple factors were recruited through the same underlying TF-family binding site. In 29% (6/21) motifs differed between the Ref and Alt alleles, suggesting allele-dependent switching between TF-family binding sites. the final 14% (3/21) showed different motif clusters for different factors on the same allele, suggesting that multiple TF/COF complexes may bind the same allele at that locus.

**Figure 3.**
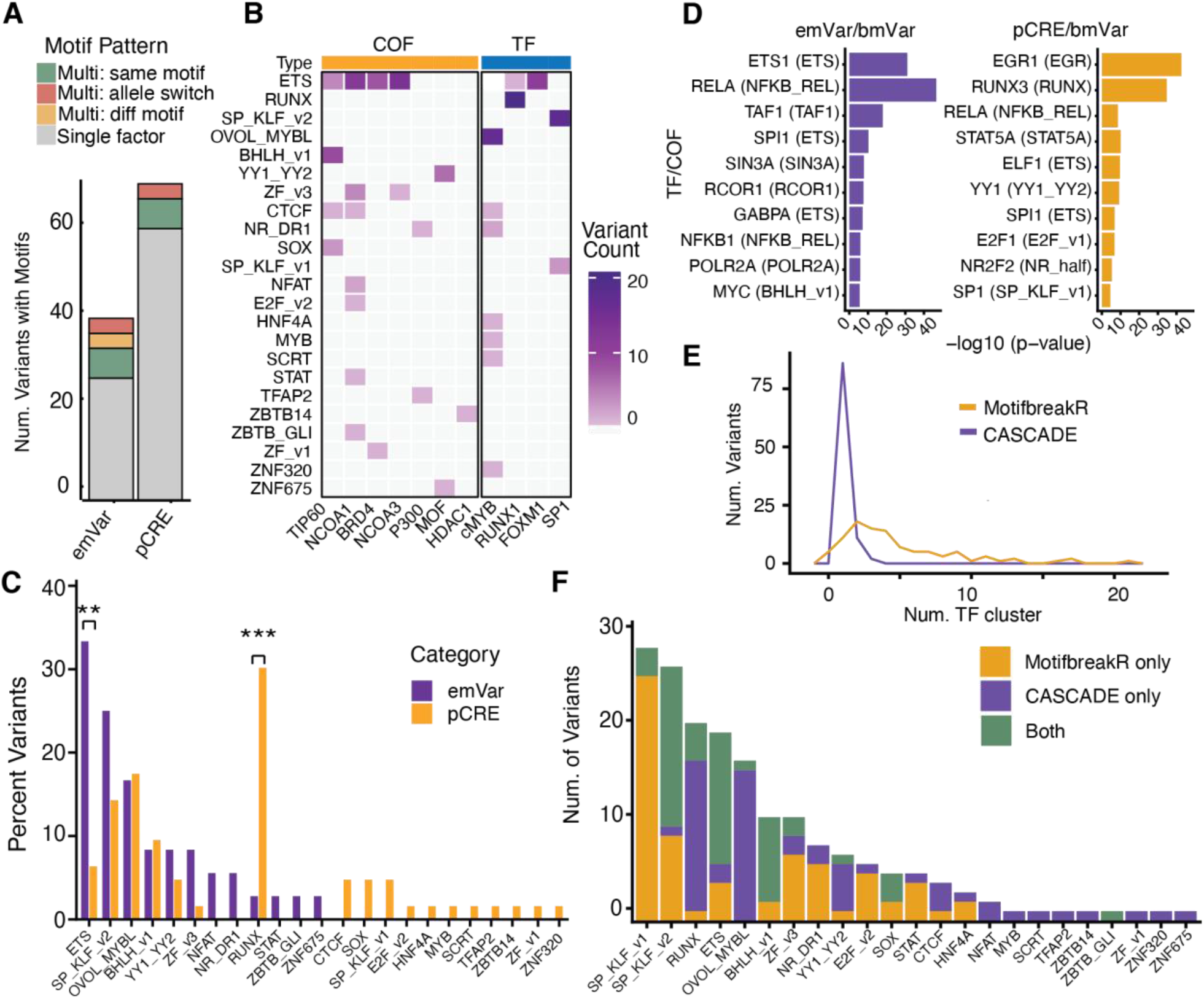
CASCADE motif analysis identifies dominant TF-family mechanisms and provides more specific regulator annotation than motifbreakR. **(a)** CASCADE Step 2 motif discovery outcomes at 99 bmVar loci, categorized as single-factor motifs, multi-factor motifs with the same motif, allele-switching motifs, or different motifs for different factors for the same allele. **(b)** CASCADE-identified motif clusters across profiled TFs and COFs. **(c)** Percentage of emVars and pCREs assigned to each motif cluster. Significance assessed by Fisher’s exact test; **p < 0.01, ***p < 0.001. **(d)** RELI enrichment of ChIP-seq datasets overlapping emVar/bmVar and pCRE/bmVar loci. **(e)** Number of TF-family clusters assigned per variant by CASCADE and motifbreakR. **(f)** Comparison of motif clusters identified by CASCADE, motifbreakR, or both methods across variants with CASCADE-determined motifs.

Across all 99 loci with one or more identified motifs, we observed 23 distinct motif clusters representing different TF families (Fig. 3b). These motif clusters are among the top clusters found enriched in putative T-cell regulatory elements defined by DHS, ATAC and H3K27ac marks in CD4+ T cells (Supplementary Figure 1), supporting their roles in T cell gene regulation, five family clusters, ETS, RUNX, SP_KLF_v2, OVOL_MYBL, and BHLH, accounted for 76% (114/150) of cluster assignments for all identified motifs. At least one TF from each of these motif clusters is expressed in Jurkat T cells (Supplementary Figure. 2), consistent with their established roles as regulators of T cell biology, and members of all five clusters have been implicated in contributing to autoimmune disease^54–56^ supporting the disease relevance of these regulators. To further assess the functional relevance of these TF families in T cells, we leveraged genome-wide Perturb-seq data from primary CD4+ T cells^57^. For each family with sufficient representation in the dataset (ETS, 16 members; SP/KLF_v1, 7 members; SP/KLF_v2, 9 members; BHLH_v1, 13 members; MYB, 1 member), we collected the union of differentially expressed genes (FDR < 10%) across all successfully knocked-down family members in resting T cells and performed GO Biological Process enrichment [Method]. Four of the five families showed significant enrichment for T cell relevant pathways including T-cell activation, adaptive immune response, leukocyte differentiation, antigen receptor signaling, T-cell differentiation, and MAPK/ERK signaling (Supplementary Figure. 3). These results confirm that the TF families with perturbed binding at autoimmune variant loci are enriched for factors that regulate T cell-specific functions.

### ETS and RUNX factors are perturbed by different classes of NCVs

To further probe variant mechanisms, we compared TF identity across variant functional classes. Surprisingly, we found different motifs were enriched at bmVars overlapping with emVars and pCREs. EmVar/bmVars were enriched for perturbed binding of ETS factors (33% vs. 6% in pCREs, p = 0.001, Fisher’s exact test) while pCRE/bmVars were enriched for perturbed binding of RUNX factors (30% vs. 3%, p < 0.001; Fig. 3c). We further examined these differences using the RELI (Regulatory Element Locus Intersection)^58^ program to systematically intersect emVar/bmVars and pCRE/bmVars with 1,544 published ChIP-seq datasets. Consistent with our CASCADE analysis, ETS family factors, including ETS1, SPI1, and GABPA, were among the most significantly enriched ChIP-seq datasets at emVar/bmVar loci, while RUNX ChIP-seq peaks showed preferential enrichment at pCRE/bmVar loci (Fig. 3d). These results highlight potential preferences of episomal MPRAs to detect certain classes of regulator variants. Consistent with the ETS enrichment we observe for emVar/bmVars, episomal MPRAs across diverse cell types consistently identify ETS-family TFs as dominant drivers of reporter gene activity, likely reflecting their broad ability to drive gene expression in the context of promoters^12,50,59–62^. In contrast, the enrichment of RUNX binding at pCRE/bmVars (Fig. 3c, d) suggests that these variants may function by mechanisms not readily captured by MPRAs. The specific enrichment of ETS and RUNX motifs at bmVars is of note as they have been identified in previous eQTL and caQTL studies in T cells. CaQTL analysis across human immune cells found that RUNX motifs were enriched among T and NK cell caQTLs while ETS motifs were enriched broadly across all immune cell types^33^. Similarly, Mu et al. identified ETS and RUNX factor motifs as the two most enriched regulator families of T-cell caQTLs^32^. Mu et al. further showed that RUNX motifs were enriched specifically among caQTLs that colocalized with eQTLs in CD8 T cells but not in monocytes, while ETS motifs were enriched among caQTLs colocalizing with eQTLs broadly across cell types, reflecting the restricted expression of RUNX factors in T/NK cells compared to the broad activity of ETS factors across immune lineages. Our results are a biophysical confirmation of these motif-based analyses of caQTL/eQTLs showing the central role of ETS and RUNX factors at autoimmune disease-associated variants.

### CASCADE provides cell-specific annotations of TF-binding not captured by motif-based methods

PWM-based approaches are widely used to annotate regulators perturbed by NCVs^63^. To evaluate whether our cell-specific CASCADE results are captured by PWM scores, we analyzed motifbreakR results^37^ for 99 variants with CASCADE-determined motifs. MotifbreakR assigned a median of four disrupted TF motif clusters per variant, whereas CASCADE assigned a median of one per variant (Wilcoxon rank-sum test, p = 2.93 × 10^−22^; Fig. 3e). The broader nomination from motifbreakR likely reflects limitations of PWMs or inclusion of TFs not present in Jurkat T cells. We next asked whether the motifbreakR-predicted TF families matched the CASCADE-identified motif clusters. MotifbreakR predicted a change in TF binding at 94/99 loci (94.9%); however, the prediction matched the CASCADE-identified motif cluster in only 51 (54.3%) of cases, while 43 (45.7%) predicted changes in different TF families (Fig. 3f). Comparing motifbreakR predictions across bmVar/emVar and bmVar/pCRE loci classes, we observed both higher prediction rate (100% vs 92.1%, respectively) and greater cluster-level agreement (64.9% vs 44.8%, respectively) for bmVar/emVars. This difference was driven in part by the differences for ETS and RUNX: motifbreakR confirmed 87.5% of CASCADE-identified ETS loci, consistent with ETS factors binding canonical, high-affinity sequences well-captured by PWM models, but confirmed only 20% of CASCADE-identified RUNX loci where TF binding occurs at sites that scored more poorly by PWMs. MotifbreakR also performed poorly on identifying disruption of OVOL/MYBL-type motifs (Fig. 3f), again suggesting that binding may occur to lower-affinity sites not captured well by motif-based methods. Together, these results show that while motifbreakR provides a sensitive sequence-based nomination of potential TF disruptions, CASCADE provides a direct biochemical readout of TF disruptions that accounts for cellular context and may capture lower-affinity motifs not captured by PWM.

### Allelic switches in TF binding sites reveal regulatory complex changes at disease-associated loci

Beyond gain or loss of TF binding, our analysis revealed more complicated situations where the variant allele disrupts the binding site for one TF while creating a binding site for a different TF. We highlight several examples of this ‘allelic switching’ that demonstrate examples of variant-induced swapping of activators and repressors. The variant rs2145623 is associated with psoriasis and IBD and is an eQTL in blood for NFKBIA^64^. NFKBIA encodes IκBα, a key inhibitor of NF-κB signaling that regulates inflammatory and immune responses. The variant is located ∼34.7 kb downstream of the NFKBIA TSS within a DHS site present in T cells, indicating that it is located in accessible chromatin in a putative regulatory region (Fig. 4a). We found that the Ref allele supports recruitment of the factors FOXM1, TIP60, BRD4, NCOA3/SRC3 and NCOA1/SRC1 to an ETS-family motif, while the alternative allele abolishes ETS-mediated recruitment and enables binding of cMYB and P300 to a newly created MYBL motif, an ETS-to-MYB regulatory switch (Fig. 4b). Although P300 is generally considered a coactivator, MYB:P300 interaction has been shown to mediate transcriptional repression at approximately half of MYB’s direct target genes^65^. MPRA data showed loss of enhancer activity with the alternative allele, consistent with either loss of the activating ETS-driven coactivator complex, a repressive function of the newly recruited MYB:P300 complex, or both. To distinguish between these possibilities, we examined caQTL data in CD4+ T cells^32^ that associates genetic variants with chromatin changes. Rs2145623 is associated with decreased chromatin accessibility (β = −0.91, adjusted p = 3.2 × 10^−8^) (Fig. 4a, c), consistent with loss of recruitment of the cofactors BRD4, TIP60, NCOA1/SRC1 and NCOA3/SRC3 that are all associated with chromatin opening. These results suggest a possible mechanism for rs2145623-dependent modulation of NFKBIA gene expression via an activator-to-repressor complex switch.

**Figure 4.**
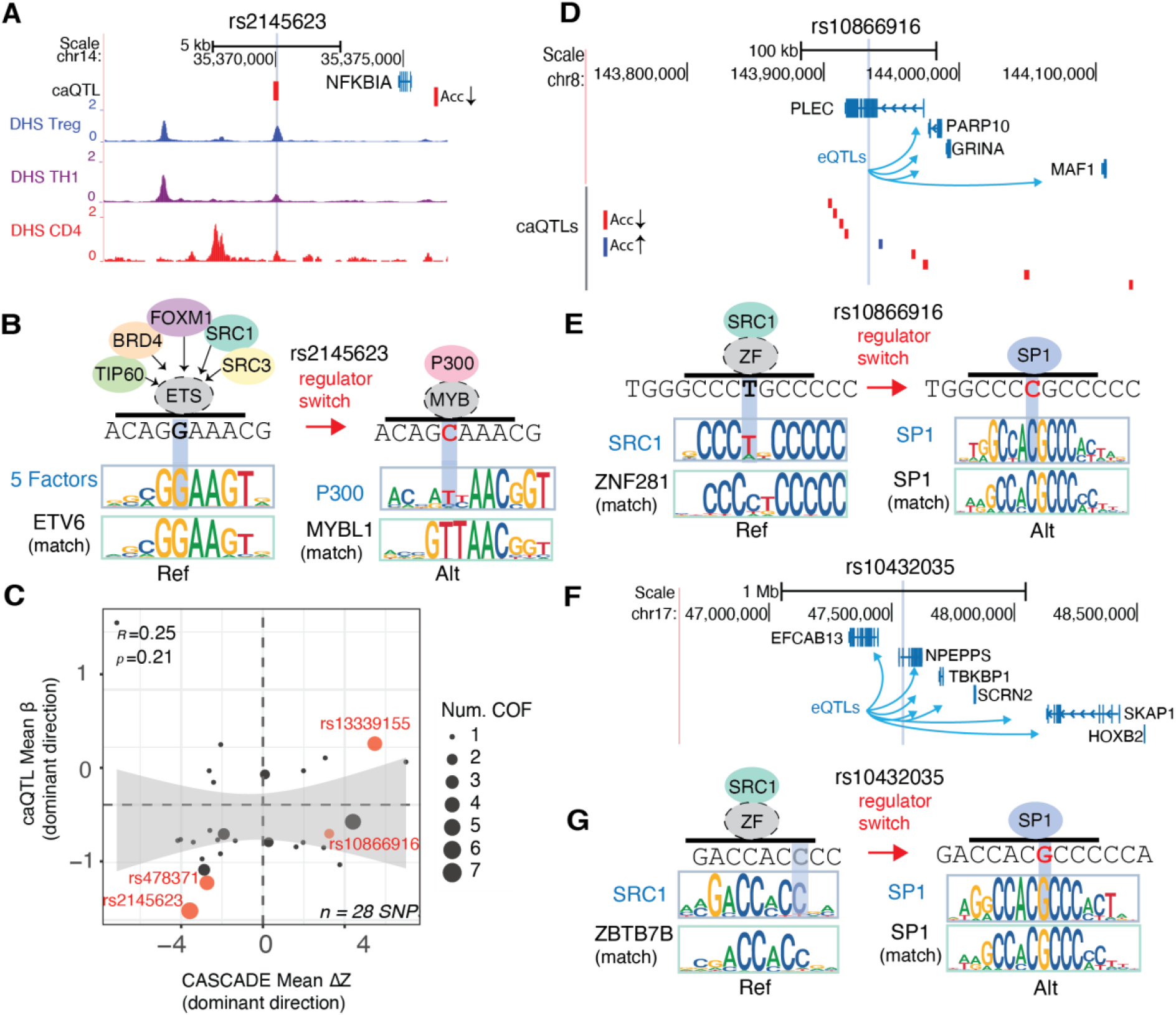
Disease-associated variants can switch TF binding sites and alter regulatory complex recruitment. **(a)** Genomic view of the rs2145623 locus near NFKBIA, showing T-cell DHS tracks and the associated caQTL accessibility effect. **(b)** Schematic of allele-dependent TF/COF recruitment showing the reference G allele and alternative C allele. A representative CASCADE motif from one profiled factor is shown for each allele (all factors showed similar motifs); a top-matched JASPAR motif is shown below. **(c)** CASCADE mean Δz-score (dominant binding direction) versus caQTL mean β (dominant accessibility direction) across 28 bmVar/emVars with caQTL data. Point size indicates the number of COFs showing differential binding. **(d)** Genomic view of the rs10866916 locus near PLEC, showing eQTL-linked genes and associated caQTL accessibility peaks. **(e)** Schematic of allele-dependent TF/COF recruitment at rs10866916, showing the reference T allele and alternative C allele. A representative CASCADE motif from one profiled factor is shown for each allele; matched JASPAR motifs are shown below. **(f)** Genomic view of the rs10432035 locus near NPEPPS, showing eQTL-linked genes. **(g)** Schematic of allele-dependent TF/COF recruitment at rs10432035, showing the reference G allele and alternative C allele. A representative CASCADE motif from one profiled factor is shown for each allele; matched JASPAR motifs are shown below.

We identified two MS-associated variants, rs10866916 and rs10432035, at distinct genomic loci that function as eQTLs in CD4+ T cells and blood, increase gene expression in MPRA, and exhibit zinc finger (ZF)-to-SP1 regulator switches. rs10866916 is located in an intron of PLEC and is associated with expression of PLEC, GRINA, PARP10, and MAF1 in CD4+ T cells and blood^64^. PLEC is involved in T cell cytoskeletal organization and chemotactic migration^66,67^ (Fig. 4d) and published promoter-capture Hi-C data in Jurkat cells^68^ confirms physical contact between the variant-containing region and the PLEC promoter. In our analysis the Ref allele supports recruitment of NCOA1/SRC1 to a ZF281 motif, while the alternative allele disrupts this binding and creates an SP1-family motif (Fig. 4e). SP1 functions as a potent activator in T cells^69^, and ZF281 has been shown to act as a repressor in CD4+ T cells^70^, consistent with the increased MPRA activity of the Alt allele. Previous caQTL analysis in CD4+ T cells^32^ revealed that rs10866916 was associated with altered chromatin accessibility at nine loci. The closest locus, located 1,400 bp upstream of the SNP, showed increased accessibility with the Alt allele (β = 0.10, adjusted p = 2.07 × 10^−4^), consistent with the MPRA gain of activity and putative SP factor-driven chromatin opening. In contrast, the other eight loci (Fig. 4d), located 2.5–33.8 kb from the variant, show decreased accessibility with the Alt allele (Fig. 4c), and eQTL analysis for the Alt allele in CD4+ T cells and blood cells^64^ show reduced expression of PLEC and other linked genes. Thus, rs10866916 illustrates a putative repressor-to-activator switch that tracks with reporter gene activity and increased accessibility at a nearby loci, while caQTL effects at more distal elements and eQTL effects in immune cells suggest a more complex locus-level regulatory outcome that may result other variants in LD.

The second ZF-to-SP1 regulator switch variant, rs10432035, is located in an intron of NPEPPS and is an eQTL for NPEPPS and other nearby genes in CD4+ T cells and blood^64^ (Fig. 4f). Our analysis reveals that the Ref allele supports recruitment of NCOA1/SRC1 to a ZBTB7B-type motif while the alternative allele creates an SP1-family motif (Fig. 4g; Supplementary Table 2). ZBTB7B/ThPOK is a known repressor in CD4+ T cells that represses CD8+ lineage genes^71,72^ and SP1, as discussed, is an activator, consistent with the increased MPRA activity of the Alt allele. However, endogenous eQTL effects at this locus are gene-, dataset-, and context-specific: some signals support increased expression of NPEPPS or nearby genes, whereas others support decreased expression of NPEPPS or other linked genes. Together these results demonstrate that NCVs can involve switches in regulator binding that may need to be accounted for when defining variant mechanisms. Further, these examples highlight that variants can cause local changes that are consistent across CASCADE and MPRA assays, while endogenous effects can diverge and may be affected by LD structure, target-gene context, and cellular state.

### Multiple NCVs at ETS motifs alter recruitment of a common suite of factors

Examining NCVs for shared mechanisms, we identified a group of seven emVar/bmVars (rs13339155, rs17346550, rs113170867, rs4672505, rs4609899, rs62054825, rs478371) that exhibited coordinated changes in recruitment for a set of shared regulators, including the TF FOXM1 and cofactors TIP60, BRD4, NCOA3/SRC3, and NCOA1/SRC1. Further, binding changes for these factors track perfectly with corresponding changes in MPRA-measured gene expression (Fig. 5a). CASCADE motif analysis identified an ETS-family motif at each of these loci (for all factors) (Fig 5b, Supplementary Table 2), indicating that ETS-family TFs recruit these factors, directly or indirectly, to activate gene expression at these loci. The fact that FOXM1 CASCADE experiments lead to a perfect ETS motif at these loci indicates indirect recruitment of the TF FOXM1 by ETS factors (Fig 5b). In this context, we hypothesize that the TF FOXM1 may function like a recruited cofactor to augment the function of the DNA-bound ETS proteins, as has been shown for FOXM1 in the regulation of cell-cycle genes^73^. The consistent co-occurrence of these five factors across seven independent loci suggests a recurring regulatory module in T cells, wherein allelic changes to an ETS motif leads to coordinated loss or gain of the entire cofactor ensemble and corresponding changes in enhancer activity. We highlight several loci to illustrate how this shared suite of regulators may operate across distinct genomic and disease contexts.

**Figure 5.**
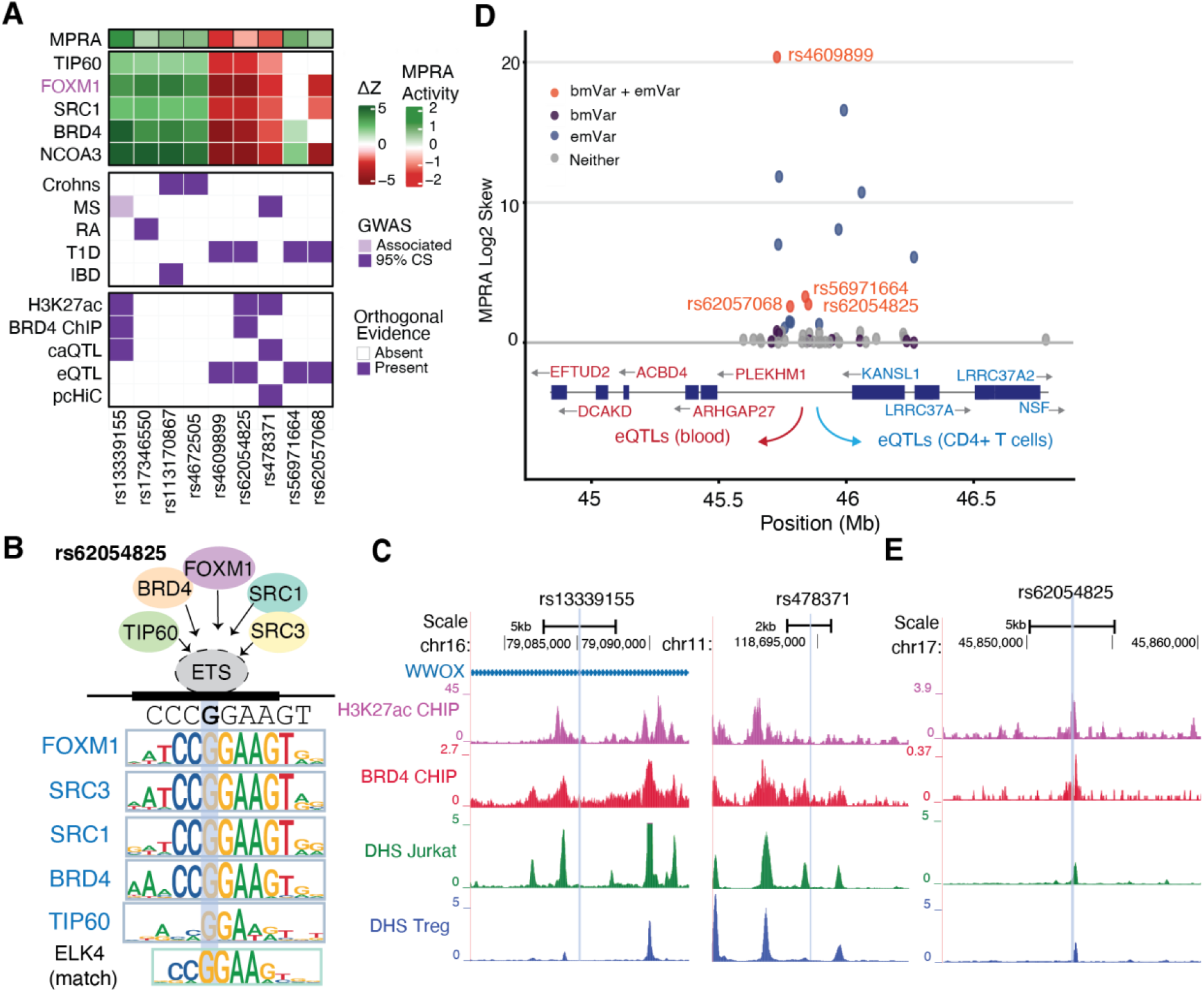
Multiple autoimmune disease-associated NCVs at ETS motifs alter recruitment of a shared regulatory module. **(a)** Summary of ETS-associated bmVar/emVars showing MPRA activity, CASCADE Δz scores for FOXM1 and cofactors, GWAS disease annotations, and orthogonal evidence including H3K27ac, BRD4 ChIP-seq, caQTL, eQTL, and pcHi-C support. The first seven variants represent the recurrent ETS-associated module; rs56971664 and rs62057068 are additional ETS-associated emVar/bmVars at the 17q21.31 locus. **(b)** Schematic of the shared ETS-associated regulatory module. CASCADE-derived motifs for the indicated factors are shown, with the matched JASPAR ETS motif shown below. **(c)** Genome browser views of rs13339155 and rs478371 showing H3K27ac, BRD4 ChIP-seq, together with nearby DHS signal in T-cell datasets. **(d)** Overview of the 17q21.31 locus showing MPRA activity across tested variants, variant regulatory classes, and nearby eQTL-linked genes in blood and CD4+ T cells. **(e)** Genome browser view of rs62054825 showing overlap with H3K27ac, BRD4 ChIP-seq, and DHS signal.

Two of the ETS-perturbing variants, rs13339155 and rs478371, are found at distinct genomic locations, are both associated with MS, coincide with caQTLs in CD4+ T cells, and overlap H3K27ac ChIP-seq peaks in T cells (Fig. 4c, 5a, Supplementary Table 5). Rs13339155 is located within an intron of *WWOX* (Fig 5c; left panel), which encodes a WW domain-containing oxidoreductase implicated in neuronal differentiation, CNS myelination, and MS susceptibility^8^. CaQTL analysis in CD4+ T cells indicates that the alternative allele is associated with increased chromatin accessibility (most significant association: β = 0.53, adjusted p = 1.12 × 10^−5^), consistent with the observed positive direction of binding and expression changes in our CASCADE and MPRA data (Fig. 5a). Rs478371 is also associated with MS (Fig 5c; right panel) and promoter-capture Hi-C data indicates that the region surrounding rs478371 physically interacts with the promoter of BCL9L, involved in T cell differentiation and immune regulation^74^. We identify a composite ETS motif at this locus rather than a canonical single ETS motif, a motif architecture commonly found in promoter elements^75^. CaQTL analysis in CD4+ T cells, showed that this variant is associated with chromatin accessibility at 7 peaks, with the majority (5/7) showing decreased accessibility for the ALT allele (most significant association: β = −0.73, adjusted p = 5.42 × 10^−16^), consistent with the direction of binding and expression changes observed in our CASCADE and MPRA data (Fig. 5a).

The 17q21.31 locus contains 50 NCVs in tight LD (r^2^ > 0.8) that were identified as associated with T1D (Fig. 5d). This region has been primarily associated with neurodegenerative and neurodevelopmental disorders, including Parkinson’s disease^76^ and Koolen-de Vries syndrome^77^, though recent analyses have also identified shared genetic architecture between this locus and T1D^78^. 43 are eQTLs for NSF, KANSL1, LRRC37A, and LRRC37A2 in CD4+ T cells^64^, four of which we identified as joint bmVar/emVars that alter ETS motifs (Fig. 5d). Two of these bmVar/emVars (rs62054825, rs4609899) belong to our set of ETS motifs found to recruit FOXM1/TIP60/BRD4/NCOA3/NCOA1 with the Alt disease-associated allele leading to a loss of ETS binding and cofactor recruitment (Fig. 5a). Among these, rs62054825 overlaps BRD4 and H3K27ac ChIP-seq peaks, and DNase I hypersensitivity (DHS) elements found in Jurkat and Treg cells (Fig. 5e), supporting its putative roles as a variant that may impact factor binding and gene expression. Together, these results suggest a recurrent regulatory module that is perturbed by disease-associated in T cells, that is variably supported by QTL studies and chromatin maps, and that involves ETS factors recruiting the TF FOXM1 and a suite of cofactors to alter gene expression.

## DISCUSSION

In this study, we profiled allele-specific TF and COF binding across 2,901 autoimmune disease-associated non-coding variants previously characterized by MPRA in Jurkat T cells. We identified 516 bmVars, ∼10% of which exhibit allele-specific enhancer activity, with strong concordance between changes in TF/COF recruitment and MPRA-measured expression effects. Integration with chromatin annotations and fine-mapping data further demonstrated that bmVar status captures functional regulatory perturbations within open chromatin elements, supporting the biological relevance of these binding changes in T cells. Motif analysis revealed that most variants perturb binding of five major TF families — ETS, RUNX, SP/KLF, OVOL/MYBL, and bHLH — all of which have established roles in T cell biology and whose motifs are enriched in putative T cell regulatory elements. Notably, different variant classes showed distinct regulatory signatures, with ETS-associated perturbations enriched among emVar/bmVars and RUNX-associated perturbations enriched among pCRE/bmVars, suggesting that different classes of autoimmune variants may act through distinct regulatory mechanisms. Our results also demonstrate that direct biochemical measurement of TF binding and COF recruitment provides more specific annotation of perturbed regulators than motif-based computational approaches alone, particularly for context-dependent binding events. At a practical level, these findings highlight the value of profiling COF recruitment in addition to TF binding, revealing regulatory variants that would not have been detected through TF-centered analyses alone. More broadly, this work demonstrates how high-throughput biochemical profiling can complement functional genomic approaches such as MPRA, chromatin profiling, and genetic fine mapping to prioritize putative causal variants and provide mechanistic insight into how non-coding disease alleles alter gene regulation.

An important feature of this study is the integration of biochemical and functional measurements for a set of prioritized autoimmune disease variants. MPRAs can identify variants that alter enhancer/promoter activity, while CASCADE adds a mechanistic layer by measuring allele-specific TF binding and COF recruitment. Among bmVar/emVars, the strong concordance between binding and reporter-expression changes supports the idea that altered TF/COF recruitment can help to explain enhancer-activity differences for prioritized variants. At individual loci, our analysis revealed mechanisms beyond simple gain or loss of binding, including TF-family switches in which one allele disrupts one regulatory complex while creating another. These regulator switches highlight the complexity in defining variant mechanisms both because effects may result from competing (or reinforcing) effects of distinct complexes, and because the effects of an allele may depend on which TFs and COFs are available in a given immune cell state. We also identified a recurrent ETS-associated regulatory module involving FOXM1 and the cofactors TIP60, BRD4, NCOA3, and NCOA1, where coordinated changes in recruitment were associated with corresponding changes in MPRA activity across multiple loci. Recent work has shown that enhancers can be classified by distinct cofactor dependencies^48^. The convergence of seven autoimmune disease-associated variants on the same ETS-associated coactivator module suggests that COF dependency may also stratify regulatory variants into mechanistically distinct classes.

Among variants with CASCADE-identified motifs, binding effects were concentrated in a small number of TF families. ETS, RUNX, SP/KLF, OVOL/MYBL, and bHLH accounted for most motifs, and Perturb-seq analysis in primary CD4+ T cells supported the relevance of these families to T-cell regulatory programs. Further, the CASCADE-defined motifs from these families showed significant enrichment in DHS, ATAC, and H3K27ac-defined regulatory elements in CD4+ T cells. Identifying these dominant TF families is informative for future variant prioritization. Rather than screening every possible TF, targeted assays focused on a defined set of T-cell-relevant TF/COF complexes may capture many regulatory mechanisms relevant to autoimmune disease-associated loci.

A surprising observation in our study was that ETS motifs were enriched among emVar/bmVars while RUNX motifs were enriched among pCRE/bmVars, suggesting that different TF classes are captured by different functional readouts. ETS-associated variants more often produced MPRA-detectable allelic effects, consistent with strong ETS activity in reporter assays. In contrast, RUNX-associated variants were enriched among MPRA-active variants without allelic expression effects, suggesting that RUNX-dependent binding may more directly alter the chromatin state, or that its impact on gene expression may be limited to different T cell states. Indeed, caQTL studies that find enrichment of RUNX motifs in specific immune cells supports these observations^32,79^. These findings fit with growing evidence that eQTLs alone do not capture all disease-relevant regulatory effects^27^. For immune-associated GWAS loci, only about one-third colocalize with eQTLs^32^, leaving many disease-associated loci without a clear expression-based mechanism .Consistent with this gap, previous studies estimated that eQTL SNPs explain only 11–14% of trait heritability^80^, whereas SNPs in enhancer and promoter regions explain a substantially larger fraction^81^, 24–79%. These observations suggest that mapping QTLs for chromatin-level phenotypes, such as caQTLs, may help uncover mechanisms at GWAS loci that are not explained by eQTLs alone. Indeed, recent single-cell caQTL mapping in human PBMCs found that caQTLs colocalize with roughly 50% more immune-disease GWAS loci than eQTLs^32^. One possibility is that some variants first affect TF/COF binding or chromatin accessibility, while gene-expression effects appear only in specific immune-cell subtypes, after stimulation, at particular time points, through distal enhancer–promoter interactions, or in combination with other linked variants. Our study highlights that CASCADE provides a complementary approach to MPRAs, or similar expression-based methods, that may better identify variants that contribute to chromatin-level regulatory effects at disease-associated loci.

Comparison of our CASCADE approach with motif-based predictions of TF binding highlights advantages for direct profiling of regulator binding. First, motifbreakR often assigned multiple TF families to the same variant, making it difficult to identify which regulator is active in the assayed cellular context. Second, motifbreakR showed lower agreement with CASCADE at RUNX and OVOL/MYBL-associated loci, consistent with the fact that PWM-based models may miss binding to lower affinity, more degenerate binding sites These findings are consistent with recent studies highlighting that predicted motif disruption alone has deficits in their ability to distinguish functional regulatory variants from background CRE variants^12^. These observations support the need for high-throughput experimental approaches to identify motif changes that are biochemically active in the relevant cellular context.

This study has several limitations that also point to future directions. Our experiments were performed using Jurkat T-cell nuclear extracts, which provide a tractable T-cell context matched to the MPRA dataset but do not capture the full diversity of primary immune cell states involved in autoimmune disease. Future studies that profile regulator binding in primary immune cell types and stimulation conditions will be important for identifying regulatory complexes that operate in disease-relevant cellular states. The current panel of five TFs and ten COFs also represents only a small fraction of the regulatory landscape. Expanding CASCADE to larger TF and COF panels, including hundreds of additional TFs and COFs available for profiling, could reveal whether distinct classes of disease-associated variants converge on shared chromatin regulators, coactivator complexes, or repressor modules. Finally, CASCADE profiles regulatory complex assembly on DNA probes outside native chromatin, while MPRAs measure enhancer activity in a reporter context. Follow-up studies using appropriately engineered cellular models will be needed to test how allele-sensitive TF/COF complexes affect endogenous chromatin accessibility and gene expression. In the long term, larger studies of perturbed TF/COF binding at disease variants should help to prioritize disease-relevant regulators for in depth analysis. We note the success of profiling COFs is of particular interest as many COFs are enzymes and, therefore, are pharmacologically targetable, whereas TFs have historically been more challenging therapeutic targets^82,83^. This study highlights that scalable biochemical profiling, when paired with functional assays, can help connect disease-associated NCVs to altered regulatory TF/COF complex assembly and allele-specific regulatory activity.

## METHODS

### Cell Culture and Nuclear Extract Preparation

Jurkat T cells were obtained from ATCC (TIB-152) and grown in suspension in RPMI 1640 Glutamax media (Thermo Fisher Scientific, Cat #72400120) with 10% heat-inactivated fetal bovine serum in T175 non-treated flasks (Thermo Fisher Scientific, Cat #132903). Nuclear extracts were prepared as previously described^49,53^. Cells were pelleted at 500×g for 5 min at 4°C and washed with 1X PBS containing protease inhibitor and pelleted again at 500 × g for 2 min at 4 °C. Plasma membranes were lysed by resuspending in 1 mL Buffer A (10 mM HEPES, pH 7.9, 1.5 mM MgCl, 10 mM KCl, 0.1 mM Protease Inhibitor, Phosphatase Inhibitor (Santa-Cruz Biotechnology, Catalog #sc-45044), 0.5 mM DTT (Sigma-Aldrich, Catalog #4315) for 10 min on ice, followed by addition of Igepal (0.1% final) and vortexing. Nuclei were pelleted at 500×g for 5 min at 4°C, resuspended in 100 µL Buffer C (20 mM HEPES pH 7.9, 25% glycerol, 1.5 mM MgCl_2_, 0.2 mM EDTA, 0.1 mM protease inhibitor, phosphatase inhibitor, 0.5 mM DTT, 420 mM NaCl), vortexed, and incubated for 1 h with mixing at 4°C. Nuclear extract was separated from debris by centrifugation at 21,000×g for 20 min at 4°C, flash-frozen in liquid nitrogen, and stored at −80°C.

### CASCADE Microarray Design

CASCADE experiments were performed using custom-designed microarrays (Agilent Technologies Inc, AMADID 086310 (step 1) and 086772 (step 2), 4×180K format). Microarray probe design followed the schema previously described^49^. All probes are 60 nucleotides long with the format: “GCCTAG” 5′ flank – 26-nt variable sequence – “CTAG” 3′ flank – “GTCTTGATTCGCTTGACGCTGCTG” 24-nt common primer. For each unique 26-nt variable sequence, five replicate probes were included in each orientation with respect to the glass slide (10 probes total per unique sequence). <u>Variant selection.</u> Autoimmune disease-associated variants were obtained from a previously published MPRA study that examined 18,312 variants from 531 GWAS loci across five autoimmune diseases (T1D, IBD, RA, psoriasis, MS) for allele-specific reporter expression^9^ [(Mouri et al., 2022)]. Variants with higher reporter expression than expected from plasmid prevalence were termed putative cis-regulatory elements (pCREs; n = 7,095), and variants with significant allelic expression differences were termed expression-modulating variants (emVars; n = 313). Our CASCADE library included 2,901 of these variants: 136 emVars, 2,552 pCREs lacking significant allelic effects, 137 inactive elements (in LD with GWAS SNPs but no MPRA activity), 50 MPRA-negative controls (250–1,000 bp from lead eQTL associations, r^2^ ≤ 0.25, no eQTL signal in Geuvadis or GTEx), and 26 CASCADE-positive controls previously shown to alter ETS-family binding^50^. <u>Design for step 1: Profiling Ref/Alt impact.</u> This microarray was designed to profile the impact of ncSNPs on TF/COF binding by comparing binding to reference (Ref) and alternate (Alt) probes. The design includes 2,901 Ref/Alt variant pairs described above, along with 2610 genomic background probes used for normalization. Each variant was represented in three registers along the microarray probe (centered, offset ±5 nt), requiring 60 individual probes on our array (3 registers x 10 replicates x 2 variants). <u>Design for step 2: Determining TF/COF binding motifs at prioritized loci.</u> This microarray was designed to determine TF or COF recruitment motifs at selected loci using the exhaustive single-variant (SV) mutagenesis approach previously described^49^. The design included probes for 56 emVar/bmVars and 113 pCRE/bmVars. For each locus, the Ref or Alt allele in the register and orientation that yielded the strongest COF binding in the Design 1 screen was selected as the starting reference sequence (seed sequence), along with all SV probes covering the 26-nt genomic locus. Using this design, each locus was characterized using 395 individual probes: (1 seed + 3 variants × 26 positions) × 5 replicates.

### CASCADE PBM Experimental Methods

CASCADE PBM experiments were performed as previously described^49,51^ using the 4×180K Agilent microarray platform. Microarrays were double stranded as described in Berger et al^84^. (2009). The double-stranded microarray was pre-wetted in HBS + TX-100 (20 mM HEPES, 150 mM NaCl, 0.01% Triton X-100) for 5 min and de-wetted in an HBS bath. Each chamber was incubated with 180 µL of nuclear extract binding mixture for 1 h in the dark, containing: 400– 600 µg Jurkat nuclear extract, 20 mM HEPES pH 7.9, 100 mM NaCl, 1 mM DTT, 0.2 mg/mL BSA, 0.02% Triton X-100, and 0.4 mg/mL salmon testes DNA (Sigma-Aldrich, Cat #D7656). The array was rinsed in HBS containing 0.1% Tween-20 and de-wetted in HBS. Arrays were then incubated for 20 min in the dark with 10 µg/mL primary antibody for the TF or COF of interest, diluted in 180 µL of 2% milk in HBS. After rinsing in HBS/0.1% Tween-20 and de-wetting, arrays were incubated for 20 min in the dark with 10 µg/mL species-matched Alexa Fluor 488- or Alexa Fluor 647-conjugated secondary antibody in 180 µL of 2% milk in HBS. Excess antibody was removed by washing twice for 3 min in 0.05% Tween-20 in HBS and once for 2 min in HBS. All primary and secondary antibodies used are listed in Supplementary Table 1. Microarrays were scanned with a GenePix 4400A scanner, fluorescence was quantified using GenePix Pro 7.2, and data were normalized using MicroArray LINEar Regression^84^. All experiments were performed in at least two replicates.

### CASCADE Computational Analysis

Image analysis and spatial detrending of microarray fluorescence intensities were performed as previously described^84^. Probe fluorescence values were transformed to z-scores using the fluorescence distribution of background probes included on each microarray. <u>Analysis for step 1: Profiling Ref/Alt impact.</u> To identify variants that alter TF/COF binding, fluorescence signals were compared between Ref and Alt alleles. Each variant was tested in six sequence arrangements (three positional registers × two strand orientations). For each arrangement, a two-sided Student’s t-test was used to compare the five replicates of the Ref allele versus the five replicates of the alternate allele. The six individual p-values were combined using Fisher’s method to generate a single aggregate p-value for each variant. The magnitude of differential binding was quantified using a Δz-score, defined as the mean z-score of alternative probes minus the mean z-score of reference probes across all registers, orientations, and replicates. To correct for testing thousands of variants, Benjamini–Hochberg correction was applied separately for each experiment, converting p-values to q-values (false discovery rate). Variants were classified as binding-modulating variants (bmVars) if they met four criteria: (1) at least one allele showed binding above background (z-score ≥ 1.0); (2) the binding difference between alleles was meaningful (|Δz| ≥ 1.0); (3) the binding difference was statistically significant (−log_10_ (p) > 3.0); and (4) the result remained significant after multiple testing correction (−log_10_ (q) > 1.3). Only variants meeting all criteria in both replicate experiments were called bmVar. Variants with detectable binding but no significant differential binding (non-bmVar allele binding) were defined by: Ref allele z-score ≥ 1.0, Alt allele z-score ≥ 1.0, −log_10_ (p) < 3.0, and −log_10_ (q) < 1.3. BmVars were further classified as “gain” (positive Δz), “loss” (negative Δz), or “mixed effect” (different factors showing opposite directions). Multi-factor support was defined as significant differential binding by two or more profiled factors, and directional concordance was assessed by determining whether all supporting factors showed the same perturbation direction. <u>Analysis for step 2: Determining TF/COF binding motifs at prioritized loci.</u> TF binding and COF recruitment motifs were determined by evaluating z-scores for seed and SV probes as previously described^49^. Δz-score matrices were calculated at each position along the probe as the difference from the median z-score across all four nucleotides at that position. To compare against known TF motifs, z-scores matrices were converted to position probability matrices (PPMs) using a Boltzmann-type transformation^49^. PPMs were compared against JASPAR 2022 motif database^85^ using TomTom (Euclidean distance, min_overlap = 6)^86^. Motifs were assigned to motif clusters based on shared binding specificity as previously described^53^.

### bmVar–emVar Concordance Analysis

To assess whether TF/COF binding changes at bmVar/emVars were directionally consistent with MPRA expression changes, we defined concordance as gain of binding (positive Δz) paired with increased MPRA expression (Log2Skew > 0), or loss of binding (negative Δz) paired with decreased MPRA expression (Log2Skew < 0). Variants with mixed binding effects, where different factors showed opposite binding directions, were excluded from concordance analyses (7.3%; 38/516 bmVars). To test whether the observed concordance rate among bmVar/emVars (77.8%) exceeded random expectation, we performed a permutation test (10,000 permutations). In each permutation, expression direction labels (Up/Down) were randomly shuffled among the bmVar/emVars while binding directions were held fixed, and concordance was recalculated. The empirical p-value was computed as the fraction of permutations with a concordance rate equal to or greater than the observed value. To test whether bmVar/emVars showed higher concordance than other variant categories, we calculated concordance rates for bmVar/pCREs and bmVar/inactive elements using the same gain/up and loss/down definition. Variants with mixed binding effects were excluded from each group, and the remaining variants were classified as concordant or discordant. BmVar/emVars were compared with each control group using a 2 × 2 contingency table containing the number of concordant and discordant variants in each group. Significance was assessed using a two-sided Fisher’s exact test.

### ChIP-seq Data Analysis

Publicly available ChIP-seq data in Jurkat T cells were compiled for TFs (ETS1, RUNX1, ELF1, MYB, GATA3, TAL1, TCF12/HEB, TCF3/E2A, LMO1), COFs (CBP, BRD4, MED1, CDK7), histone marks (H3K27ac, H3K27me3, H3K4me3), and RNA Pol II from multiple published studies (GEO accession numbers listed in Supplementary Table 5). Where pre-called peaks were available, peak files were downloaded directly. Where only raw sequencing reads were available, FASTQ files were downloaded from SRA, aligned to the hg38 reference genome using Bowtie2, and processed through a standard pipeline: SAM-to-BAM conversion, sorting, and indexing with SAMtools (v1.12), followed by PCR duplicate removal with Picard. Peaks were called using MACS2 with both narrow and broad peak modes (q-value threshold = 0.05), with input controls where available. Datasets originally mapped to hg18 or hg19 were converted to hg38 coordinates using UCSC liftOver. Signal tracks (bigWig format) were generated from deduplicated BAM files using deepTools bamCoverage (v3.5.1) with CPM normalization and 25-bp bins; signal tracks originally in WIG format were converted to hg38 via liftOver through bedGraph intermediate format. All tracks were visualized on the UCSC Genome Browser. Overlap between ChIP-seq peaks and variant positions was determined using GenomicRanges findOverlaps in R, where each variant was represented as a single-nucleotide GRanges object and tested for overlap with peak intervals.

### PICS, DHS, and H3K27ac Enrichment Analyses

Enrichment of bmVars in chromatin features and fine-mapped variant sets was assessed using two complementary analyses. DHS overlap was determined using a merged T cell DHS peak set combining DNase-seq datasets from different T cell subtypes available in ENCODE (Supplementary Table 5). H3K27ac overlap was defined as intersection with peaks in at least two of five published Jurkat H3K27ac ChIP-seq datasets (Supplementary Table 5). Variant–peak overlaps were computed as described in the ChIP-seq Data Analysis section. PIP scores were obtained from *Mouri et al*^9^. and are provided in Supplementary Table 3, Sheet 3; for each variant, the PIP score from the disease with the strongest association was used.

To compare chromatin feature overlap across functional categories (Fig. 2d), each variant was assigned to one of five categories: bmVar/emVar (bm + emVar), non-bmVar emVar (emVar only), bmVar/pCRE (bm + pCRE), non-bmVar pCRE (pCRE only), or non-bmVar/MPRA-inactive baseline (Non bm/emVar). For each category, the fraction of variants overlapping DHS or H3K27ac was calculated. Enrichment was assessed by comparing each category to the non-bmVar/MPRA-inactive baseline using a two-sided Fisher’s exact test. To assess PIP score enrichment among bmVars (Fig. 2e), we calculated the fraction of variants with PIP scores above each threshold: 0.01, 0.05, 0.1, 0.2, 0.3, 0.4, and 0.5. For bmVar/emVar and bmVar/pCRE comparisons without chromatin filtering, fold enrichment was calculated relative to the corresponding non-bmVar category: bmVar/emVars versus non-bmVar emVars, and bmVar/pCREs versus non-bmVar pCREs. For DHS- and H3K27ac-overlap comparisons, fold enrichment was calculated as the fraction of variants in the indicated bmVar/chromatin-overlap category above each PIP threshold divided by the fraction of all other tested variants not in that category above the same threshold. Significance at each threshold was assessed using a two-sided Fisher’s exact test.

### TF Expression in Jurkat T Cells

To confirm that TFs from the top CASCADE-identified motif clusters are expressed in Jurkat T cells, we used normalized protein abundance values for Jurkat cells from published quantitative proteomics data^87^. TFs were assigned to motif clusters using published binding motif clusters assignments from our lab^53^. For genes with multiple protein isoforms, abundance values were averaged. Protein levels for TFs in each of the top five motif clusters (ETS, RUNX, SP_KLF, OVOL_MYBL, BHLH) were compared to the distributions of all detected proteins and all detected TFs in Jurkat cells, with the median protein level of housekeeping genes (ACTB, GAPDH, TUBB, B2M, HPRT1, TBP, RPLP0, PPIA) shown as a reference (Supplementary Fig. 2).

### Genome-wide Perturb-seq Analysis

Genome-wide Perturb-seq differential expression statistics in primary human CD4+ T cells were obtained from Zhu et al^57^. The DE statistics file (GWCD4i.DE_stats.h5ad) was downloaded from the associated AWS S3 bucket (s3://genome-scale-tcell-perturb-seq/marson2025_data/). This dataset contains DESeq2 results for ∼11,527 CRISPRi perturbation targets across ∼10,282 genes in three conditions (Rest, Stim8hr, Stim48hr). Using a custom Python pipeline, we extracted DE statistics for two sets of perturbation targets: (1) the 15 TFs and COFs profiled in CASCADE Step 1 (SP1, FOXM1, MYB, RUNX1, LEF1, KAT5/TIP60, EP300, NCOA1/SRC1, NCOA3/SRC3, HDAC1, BRD4, MED23, MOF, RUVBL2), of which 12 were present in the dataset (MOF and RUVBL2 were absent); and (2) some TF family members belonging to motif clusters identified in CASCADE Step 2 (ETS, RUNX, SP_KLF v1/v2, BHLH_v1, OVOL_MYBL, ZBTB_GLI). Cluster membership was defined using the Siggers lab motif cluster database^53^ (Supplementary Table4). Perturbation targets were matched using synonym-based lookup against curated TF and COF reference lists. The analysis was restricted to perturbations with confirmed successful knockdown (ontarget_significant == TRUE), with downstream effects considered significant at FDR < 10% (adjusted p-value < 0.10). For each TF family represented in the Perturb-seq dataset (ETS, MYB, BHLH_v1, SP_KLF_v1, SP_KLF_v2), GO Biological Process enrichment was performed on the union of genes significantly differentially expressed (adjusted p-value < 0.10) following knockdown of any family member in resting T cells. Enrichment was performed using gprofiler2 [REF] against a background of all genes tested across successful knockdowns in the resting condition. Enriched terms were considered significant at Benjamini–Hochberg adjusted p-value < 0.05. The RUNX family was excluded because only RUNX3 passed knockdown QC (RUNX1 and RUNX2 failed in all conditions) and yielded only 9 DE genes, too few for enrichment analysis. The OVOL_MYBL cluster was similarly excluded because only MYBL1 was present in the dataset and yielded only 2 DE genes.

### RELI Analysis

To assess enrichment of in vivo TF and COF binding at bmVar/emVar and bmVar/pCRE loci, genomic coordinates (hg19) for 36 bmVar/emVar and 63 bmVar/pCRE loci with CASCADE-identified motifs were intersected against 1,544 published ChIP-seq datasets using the RELI algorithm^58^. RELI estimates significance by generating a null distribution through 2,000 permutations of size- and minor allele frequency-matched random SNP sets drawn from dbSNP, computing a Z-score, relative risk, and empirical p-value for each dataset. P-values were Bonferroni-corrected across all 1,544 datasets. Datasets where no permutation generated any overlap (relative risk = 2,001) were excluded as permutation-based statistics are unreliable for sparse ChIP-seq peak sets. Significant enrichment was defined as Bonferroni-corrected p < 0.05. For visualization, the most significant hit per regulatory factor per variant category was retained and the top 10 enriched factors per category are shown.

### MotifbreakR Comparison Analysis

CASCADE-determined motif assignments were compared against motifbreakR predictions^37^. Predicted TFs were assigned to motif clusters using the same CoRec-based cluster definitions applied to CASCADE motifs^53^. A locus was considered concordant if any motifbreakR-predicted TF belonged to the same motif cluster as the CASCADE-assigned motif. For each variant, we counted the number of distinct TF motif clusters assigned by CASCADE and motifbreakR, and compared the distribution of per-variant cluster counts between methods using a Wilcoxon rank-sum test.

### eQTL and Promoter-Capture Hi-C Annotations

eQTL and promoter-capture Hi-C (pcHi-C) annotations were obtained from Open Targets Variant-to-Gene (V2G) data^64^ and are provided in Supplementary Table 3, Sheet 3. V2G gene assignments are based on distance to variant, eQTL, sQTL, pQTL, and pcHi-C data, filtered for genes expressed in primary T cells (TPM > 1).

### Chromatin Accessibility QTL (caQTL) Analysis

caQTL summary statistics for CD4+ T cells were obtained from Mu et al. (2026)^32^. Full summary statistics were downloaded from Zenodo (https://doi.org/10.5281/zenodo.14652965; file: full_summstats_CD4.T_250kb.txt.gz), covering 85% (2,439/2,884) of our CASCADE variants. As each variant is associated with multiple chromatin accessibility peaks, p-values (PVALUE_RE2) were adjusted using Benjamini–Hochberg correction within each peak. Variants were classified as significant caQTLs at adjusted p-value < 0.01. β coefficients and adjusted p-values reported in the Results reflect the most significantly associated peak for each variant.

### Motif enrichment analysis

Genomic locations of chromatin-accessible regions in CD4+ T cells, including DNase-seq and ATAC-seq peaks, together with H3K27ac ChIP-seq peaks, were obtained from ENCODE Project Consortium^88^ (Supplementary Table 5) and analyzed for motif enrichment using HOMER^89^. All genomic regions were formatted to be peak-centered, 500 bp-wide based on downloaded BED files. HOMER was run using a custom background region (BG). BG regions were 500-bp in length selected from the hg38 genome and both GC fraction-matched and CpG fraction-matched to the input regions. The size of the BG matched the number of foreground regions in each set. HOMER was run using the full JASPAR 2022 core set of motifs^85^. To define the enrichment of our CASCADE-identified motifs compared to all motifs in these T cell genomic elements (Supplementary Table 3, sheet 2), motifs were grouped into TF motif-classes by motif similarity (Supplementary Table 4), all classes were ranked according to maximum - log(p-value) from HOMER, and we used Wilcoxon-Mann Whitney U-statistic (visualized as an ROC curve) to calculate the enrichment of CASCADE-motifs against the full JASPAR motif database.

## Data availability

All processed data generated in this study are provided in the Supplementary Tables. Raw and processed CASCADE microarray data will be deposited in a public repository, such as GEO or ArrayExpress, before publication.

## Code availability

Analysis code used in this study will be made available upon request.

## Acknowledgements

We thank Dr. John Ray for his thoughtful comments on the manuscript and helpful discussions related to the MPRA analysis, and Drs. Zeba Wunderlich, Brian Clary, and Juan Fuxman Bass for helpful feedback and discussions. M.D. was supported by a NIH-funded predoctoral training fellowship (T32GM130546) and a Kilachand Fellowship made through the Multicellular Design Program (MDP) at Boston University. This work was supported by NIH grants (R01AI51051 and R21HG011289) to T.S.

## Author contributions

M.D. and T.S. conceived and designed the study. M.D. performed the experiments, analyzed the data, and interpreted the results with input from T.S.; M.D. and T.S. wrote the manuscript; T.S. supervised the study. Both authors reviewed and approved the final manuscript.

## Supplementary Figures

**Supplementary Figure 1.**
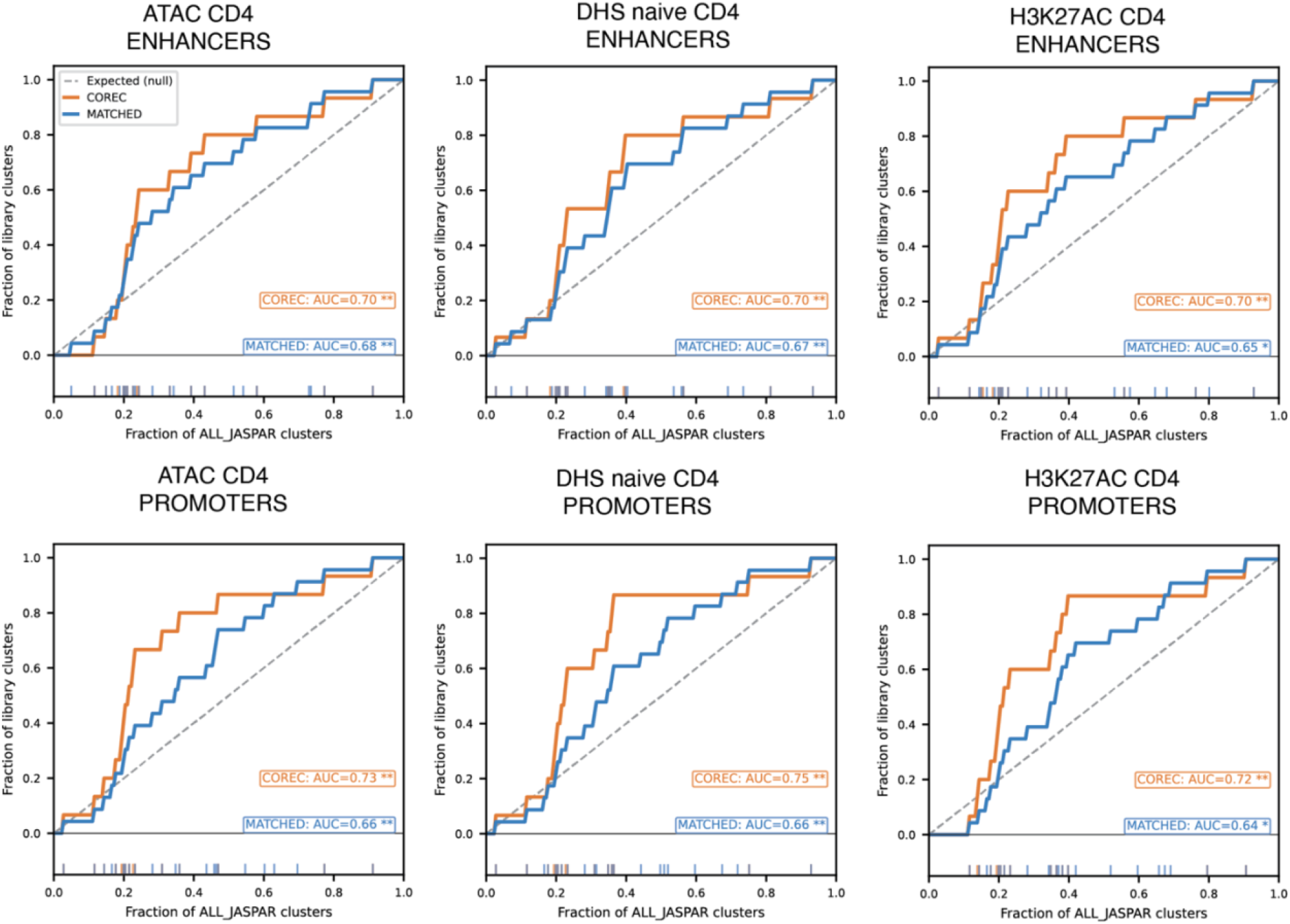
Top CASCADE motif clusters are enriched in CD4+ T-cell regulatory elements. ROC-style enrichment curves showing the rank enrichment of CoRec and matched motif clusters across CD4+ T-cell regulatory element datasets defined by ATAC-seq, naive CD4 DHS, and H3K27ac marks. Motif clusters were ranked using HOMER enrichment scores across all JASPAR clusters. The x-axis shows the cumulative fraction of all JASPAR clusters ranked by score, and the y-axis shows the cumulative fraction of library clusters seen. Separate panels show enhancer and promoter annotations. The dashed line indicates the expected null distribution. AUC values are shown for CoRec and matched motif-cluster rankings.

**Supplementary Figure 2.**
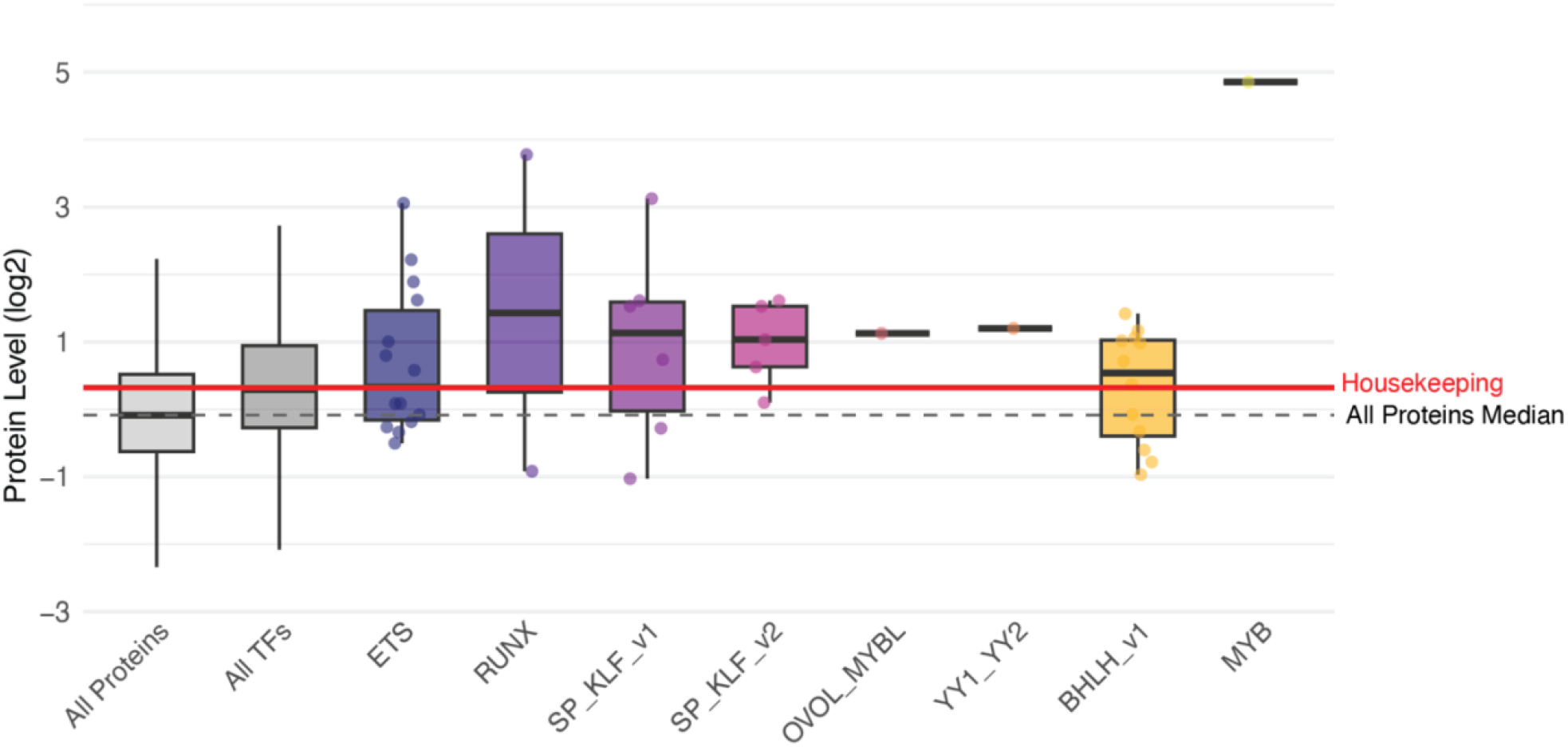
Protein abundance of TFs from top CASCADE motif clusters in Jurkat cells. Boxplots showing normalized protein abundance distributions for TFs assigned to major CASCADE-identified motif clusters in Jurkat cells, compared with all detected proteins and all detected TFs. Each overlaid point represents an individual TF member within the indicated motif cluster. Protein levels are shown on a log2 scale, and the housekeeping protein median is shown as a reference.

**Supplementary Figure 3.**
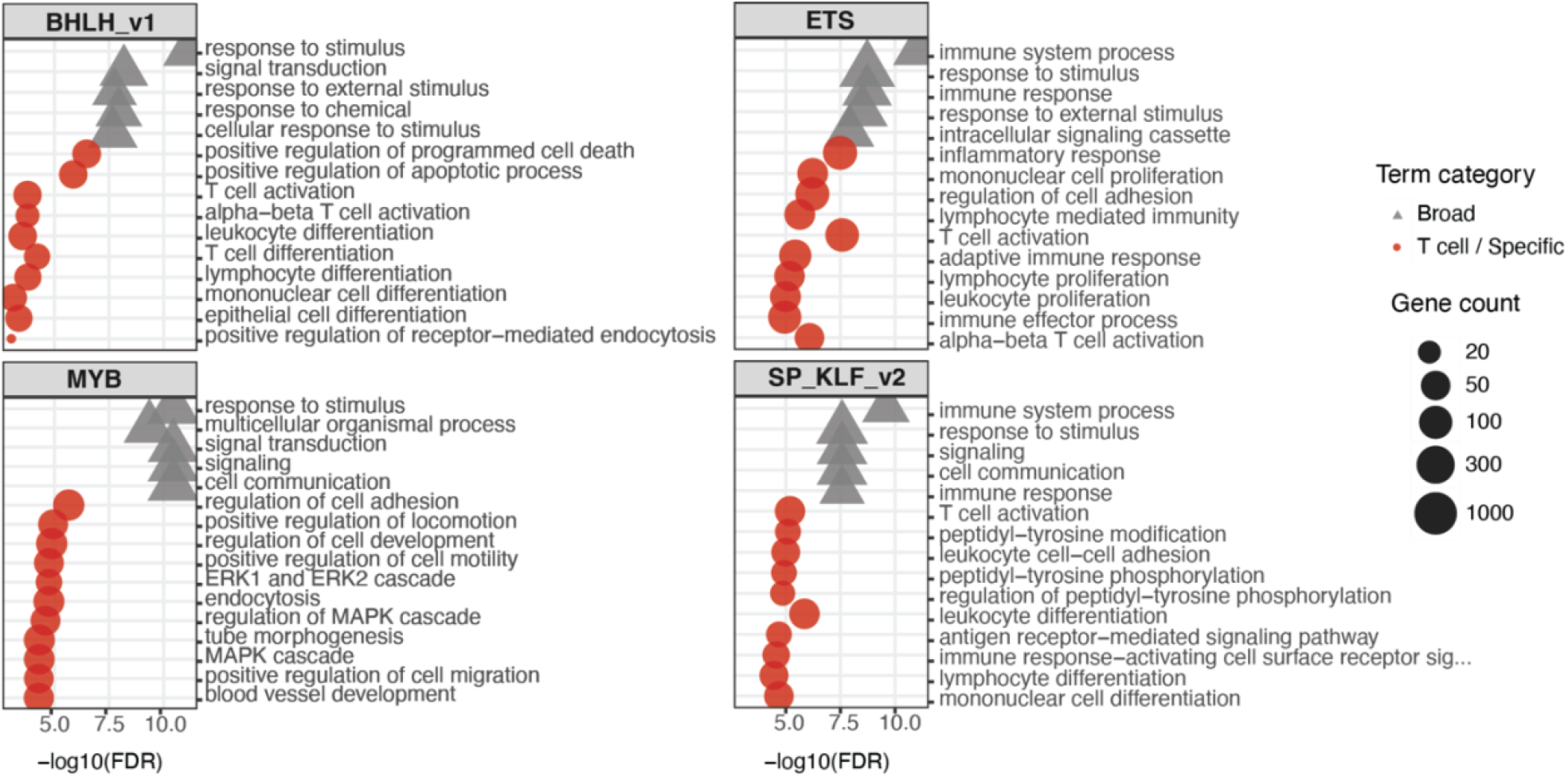
Perturb-seq GO enrichment supports T-cell regulatory roles for dominant CASCADE motif families. GO Biological Process enrichment analysis of genes differentially expressed after knockdown of TF-family members in resting primary CD4+ T cells. Enrichment results are shown for BHLH_v1, ETS, MYB, and SP_KLF_v2 families. Terms are categorized as broad biological processes or T-cell/specific immune-related processes. Point size indicates the number of genes associated with each term, and the x-axis shows −log10(FDR).

